# Mismatch repair hierarchy of *Pseudomonas putida* revealed by mutagenic ssDNA recombineering of the *pyrF* gene

**DOI:** 10.1101/710293

**Authors:** Tomas Aparicio, Akos Nyerges, István Nagy, Csaba Pal, Esteban Martínez-García, Víctor de Lorenzo

## Abstract

The mismatch repair (MMR) system is one of the key molecular devices that prokaryotic cells have for ensuring fidelity of DNA replication. While the canonical MMR of *E. coli* involves 3 proteins (encoded by *mutS, mutL and mutH*), the soil bacterium *Pseudomonads putida* has only 2 *bona fide* homologues (*mutS* and *mutL*) and the sensitivity of this abridged system to different types of mismatches is unknown. On this background, sensitivity to MMR of this bacterium was inspected through single stranded (ss) DNA recombineering of the *pyrF* gene (the prokaryotic equivalent to yeast’s URA3) with mutagenic oligos representative of every possible mispairing under either wild-type conditions, permanent deletion of *mutS* or transient loss of *mutL* activity (brought about by the thermoinducible dominant negative allele *mutL_E36K_*). Analysis of single nucleotide mutations borne by clones resistant to fluoroorotic acid (5FOA, the target of wild type PyrF) pinpointed prohibited and tolerated single-nucleotide replacements and exposed a clear grading of mismatch recognition. The resulting data unequivocally established the hierarchy A:G< C:C< G:A< C:A, A:A, G:G, T:T, T:G, A:C, C:T< G:T, T:C as the one prevalent in *Pseudomonas putida*. This information was vital for enabling recombineering strategies aimed at single-nucleotide changes in this biotechnologically important species.

**Originality-Significance Statement:** Single-stranded DNA (ssDNA) recombineering has emerged in recent years as one of the most powerful technologies of genome editing in *E. coli* and other Enterobacteria. However, the efforts to expand the concept and the methods towards environmental microorganisms such as *Pseudomonas putida* have been limited thus far by several gaps in our fundamental knowledge of how nucleotide mismatch repair (MMR) operates in such non-model species. One critical bottleneck is the hierarchy of recognition of different types of base mispairings as well as the need of setting up strategies for counteracting MMR and thus enabling tolerance to all types of changes. The work presented here tackles both issues and makes *P. putida* amenable to sophisticated genetic manipulations that were impossible before.

## INTRODUCTION

Mutations in DNA are often caused by small insertion-deletion loops generated by strand slippage during replication and/or misincorporation of bases—by themselves or damaged by oxidative stress or other modifications (Wyrzykowski and Volkert, 2003; Putnam, 2016). The resulting base pairing mismatches are most frequently fixed by mechanisms that are remarkably conserved through the prokaryotic realm (Putnam, 2016). The steps involved in such a repair involve the recognition of an unusual structure in the DNA helix caused by the mismatch, excision of the last-synthesized strand to the site of mispairing and *de novo* synthesis of the earlier excised strand. In order to avoid inheritance of mutations, the critical feature of mismatch repair (MMR) systems is distinguishing between the old, non-modified DNA strand that acts as template and the new DNA sequence bearing the lesion. In the case where the issue has been examined in more depth (*Escherichia coli*), such a discrimination seems to be feasible owing to the interplay between its MMR and the *dam* methylation system for d(GATC) sites.

According to the current model, the MMR machinery is recruited towards the strand that is transiently unmethylated after replication. This intricate process is effected through the concerted action of three proteins encoded by *mutS* (mispair recognition), *mutL* (signal propagation) and *mutH* (strand discrimination), the action of which is then followed by DNA excision and resynthesis involving additional proteins UvrD and MutU (Matson and Robertson, 2006).

Inspection of homologous genes in a variety of bacterial branches suggests many species-specific adaptations around the archetypal MMR of *E. coli*. MutS and MutL variants have been found in virtually all Gammaproteobacteria, but MutH is often missing in many other members of the group (Putnam, 2016). This raises questions on whether the same deformations of the DNA helix are recognized by the MMR systems (mostly by MutS) in all species and how DNA strand discrimination occurs in bacteria lacking *mutH*. This question has a direct consequence on the hierarchy of mismatch recognition, as each of the 12 possible mispairings (A:G, C:C, G:A, C:A, A:A, G:G, T:T, T:G, A:C, C:T, G:T and T:C) should generate a distinct type of distortion in the DNA structure. Intuitively, the mechanism just described would predict that the bulkier the mismatch is, the easier it is to detect by the MMR system and thus fixed. But it is also possible that MutS specializes in different mispairings in diverse species. As a consequence, the MMR may become blind to some nucleotide changes, which could thus be propagated into the progeny in some hosts while others would be instead quickly removed. This originates a hierarchy of mismatch recognition by MMR, which has been clearly established in *E. coli* and other enterobacteria (Kramer *et al*., 1984; Babic *et al*., 1996; Joshi and Rao, 2001; Nyerges *et al*., 2016) but it less known in most others (Long *et al*., 2014; Long *et al*., 2018). Data on such recognition order is of essence for planning recombineering experiments aimed at introducing single nucleotide changes at specific genomic sites, as they can be counteracted with various efficiencies by the native MMR system of the target species and strains (Wang *et al*., 2009; Aparicio *et al*., 2016; Nyerges *et al*., 2016).

As is the case with other *Pseudomonas, P. putida* strain KT2440 has *mutS* and *mutL*, but lacks both *mutH* and a *dam* methylation system. Unlike *E. coli*, strand discrimination in this species could occur not through methylation but possibly through a device somehow embedded in the replication machinery itself, but the state of affairs in *Pseudomonas* is uncertain at this time (Oliver *et al*., 2002; Saumaa *et al*., 2006; Tark *et al*., 2008). In view of the growing importance of *P. putida* as a platform for synthetic biology-guided metabolic engineering and the benefits of implementing high efficiency genome editing methods (i.e. MAGE; Wang, 2009 #62} and DIvERGE (Nyerges *et al*., 2018)), it became of essence to set unequivocally the recognition preference of its native MMR system for each of the possible single nucleotide mispairs.

In the work below presented below we have capitalized on the availability of a *P. putida-born*, Erf-like recombinase (called Rec2) and a simple single-stranded (ss) DNA recombineering protocol (Ricaurte *et al*., 2018) for exploring the whole recognition landscape of mismatches that can be introduced in the genomic DNA of strain EM42 of this species (Martinez-Garcia *et al*., 2014). By inspecting the distribution of single-nucleotide changes covering the whole spectrum of mispairs we authenticated the ease of replacement of given bases by any of the others in MMR-plus and MMR-minus genetic backgrounds. The outcome turned out to be similar, but not identical, to what is known for *E. coli*. The robust recombineering approach for inspecting the MMR adopted in this work has been instrumental for setting a much improved method that can be of general value for unraveling the same question—and expanding recombineering in general— to many other bacterial species.

## RESULTS AND DISCUSSION

### *A genetic platform for inspecting MMR in* P. putida

The starting point of this work is the notion that inhibiting MMR should result in the bias-free incorporation of all possible base substitutions in the DNA helix *in vivo* (Nyerges, 2016). Such an inhibition could be made permanent e.g. through deletion of *mutS*, or transient e.g. through conditional expression of a dominant negative allele of either *mutS* (Wu and Marinus, 1994) or *mutL* (Aronshtam and Marinus, 1996; Nyerges *et al*., 2016). While deleting *mutS* is straightforward with genetic methods available for *P. putida* (see *Experimental procedures*), the second scenario (temporary suppression of MMR genes) required a different strategy. Inspection of the *mutL* (PP_4896) of *P. putida* indicated a 44 % aa identity with the orthologue of *E. coli*, which was more pronounced in their N-terminal half of the corresponding proteins—where the segments important for their function lay (Ban and Yang, 1998; Putnam, 2016). A E32K change in such an N-terminal of *E. coli*’s MutL is known to generate a mutated, inactive protein that, when over expressed from a plasmid, behaves as a negative dominant allele capable of impairing the activity of the MMR machinery of *E. coli* in the presence of the chromosomal, wild-type copy (Aronshtam and Marinus, 1996; Nyerges *et al*., 2016). This location is equivalent to conserved amino acid position 36 of the *P. putida*’s homologue (Fig. 1, Supplementary Fig. S1) and therefore we reasoned that overexpression *in vivo* of the variant *mutL*_E36K_^PP^ could bring about the same effect in this species. Finally, in order to enter mismatches of different types in a target DNA sequence, we thought of exploiting the ability of the Rec2 recombinase to enable the invasion of the replication fork of *P. putida* by synthetic single-stranded (ss) oligonucleotides *in vivo* (Ricaurte *et al*., 2018).

**Figure 1.**
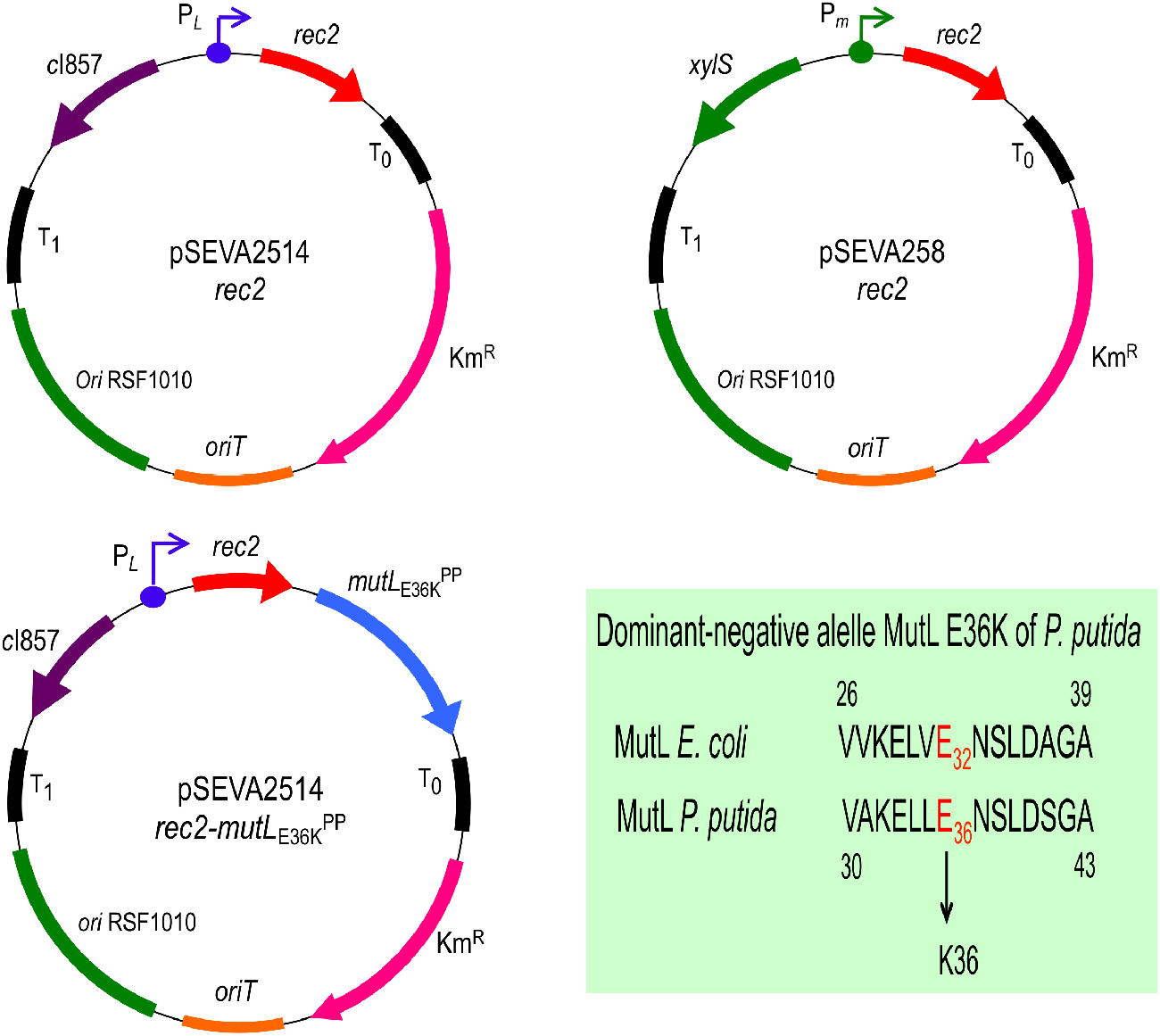
Plasmids used in this study Structure of plasmids promoting recombineering are shown(T_0_ and T_1_, transcriptional terminators; Km, Kanamycin resistance gene; *ori*T, origin of transfer; *ori* RSF1010, origin of replication; *c*I857-P_*L*_, temperature inducible expression system; *xylS*-P_*m*_, expression system inducible by 3-MB; *rec2*, recombinase; *mutL*_E36K_^PP^, dominant-negative allele of *mutL*). A conserved amino acid stretch of *E. coli* and *P. putida* KT2440 MultL proteins is also shown. The change E→K, responsible of the dominant-negative phenotype over MMR system, is highlighted in red (see Supplementary Fig. S1 for complete alignment). Pictures are not drawn to scale. pSEVA2514-*rec2* map derives from (Ricaurte *et al*., 2018).

On these bases, we generated, in one hand, an MMR-null strain by erasing *mutS* altogether (Table 1), which was used as a reference for complete elimination of mismatch repair. *P. putida* EM42 Δ*mutS* was constructed by deleting a 0.7 Kb region of the gene PP_1626 by ssDNA recombineering/CRISPR-Cas9 (see details in *Experimental Procedures*). The phenotype of this strain was tested with a rifampicin resistance (Rif^R^) assay, which was performed as a proxy of the mutational state of caused by the deletion. Results shown in Supplementary Fig. S2A accredited that—as expected— the deleted mutant underwent a much higher spontaneous mutational regime that its parental strain, which can be attributed to the loss of MMR system.

**Table 1.** Bacterial strains and plasmids used in this work.

| Strain or plasmid | Relevant characteristics <sup>a</sup> | Reference or source |
| --- | --- | --- |
| *Escherichia coli* |  |  |
| CC118 | Cloning host; $\Delta(ara-leu)$ *araD* $\Delta lacX174$ *galE* *galK* *phoA* *thiE1* *rpsE*(Sp<sup>R</sup>) *rpoB* (Rif<sup>R</sup>) *argE*(Am) *recA1* | (Manoil and Beckwith, 1985) |
| HB101 | Helper strain used for conjugation; F<sup>-</sup> $\lambda^-$ *hsdS20*(rB<sup>-</sup> mB<sup>-</sup>) *recA13* *leuB6*(Am) *araC14* $\Delta(gpt-proA)62$ *lacY1* *galK2*(Oc) *xyl-5* *mtl-1* *thiE1* *rpsL20*(Sm<sup>R</sup>) *glnX44*(AS) | (Boyer and Roulland-Dussoix, 1969) |
| *Pseudomonas putida* |  |  |
| EM42 | KT2440 derivative; Δprophage1 Δprophage4 Δprophage3 Δprophage2 ΔTn7 ΔendA-1 ΔendA-2 ΔhsdRMS Δflagellum ΔTn4652 | (Martinez-Garcia *et al.*, 2014) |
| EM42ΔmutS | EM42 derivative; ΔmutS | This work |
| Plasmids |  |  |
| pSEVA2514 | Inducible expression vector; *oriV*(RFS1010); cargo [cl857-P<sub>L</sub>]; standard multiple cloning site; Km<sup>R</sup> | (Aparicio *et al.*, 2019b) |
| pSEVA258-ssr | pSEVA258 derivative bearing the *ssr* recombinase; *oriV* (RFS1010); cargo [*xyIS-Pm* → *ssr*]; Km<sup>R</sup> | (Ricaurte *et al.*, 2018) |
| pSEVA258-ssr-*mutL*<sub>E36K</sub><sup>PP</sup> | pSEVA258 derivative bearing the *ssr* recombinase and *mutL*<sub>E36K</sub><sup>PP</sup> allele; *oriV* (RFS1010); cargo [*xyIS-Pm* → *ssr-mutL*<sub>E36K</sub><sup>PP</sup>]; Km<sup>R</sup> | This work |
| pSEVA2514-*rec2* | pSEVA2514 derivative bearing the *rec2* recombinase; *oriV*(RFS1010); cargo [cl857-P<sub>L</sub> → *rec2*]; Km<sup>R</sup> | This work |
| pSEVA2514-*rec2*-*mutL*<sub>E36K</sub><sup>PP</sup> | pSEVA2514 derivative bearing the *rec2* recombinase and *mutL*<sub>E36K</sub><sup>PP</sup> allele; *oriV*(RFS1010); cargo [*cl857*-P<sub>L</sub> → *rec2*-*mutL*<sub>E36K</sub><sup>PP</sup>]; Km<sup>R</sup> | This work |
| pSEVA231-CRISPR | pSEVA231 derivative bearing the CRISPR array; *oriV* (pBBR1); Km<sup>R</sup> | (Aparicio *et al.*, 2018) |
| pSEVA231-C-*mutS*1 | pSEVA231 derivative bearing the CRISPR array with a *mutS* spacer; *oriV* (pBBR1); Km<sup>R</sup> | This work |
| pSEVA421-Cas9tr | pSEVA421 derivative bearing the *cas9* gene and *tracrRNA*; *oriV* (RK2); Sm<sup>R</sup>/Sp<sup>R</sup> | (Aparicio *et al.*, 2018) |
| pSEVA658-*ssr* | pSEVA658 derivative bearing the *ssr* recombinase; *oriV* (RSF1010) ); cargo [*xyIS*-P<sub>m</sub> → *ssr*]; Gm<sup>R</sup> | (Aparicio *et al.*, 2018) |
| pRK600 | Helper plasmid used for conjugation; *oriV*(ColE1), RK2 (mob+ tra+); Cm<sup>R</sup> | (Kessler *et al.*, 1992) |

<sup>a</sup> Antibiotic markers: Km, kanamycin; Rif, rifampicin; Sm, streptomycin; Sp, spectinomycin; Cm, chloramphenicol; Gm, gentamicin

On the other hand, we constructed two conditional expression plasmids for either *rec2* alone or the same but assembled in the same transcriptional unit together with *mutL*_E36K_^PP^. In either case, the expression cargo (whether *rec2* alone or *rec2*-*mutL*_E36K_^PP^) was placed under the control of the heat-inducible *c*I857-P_L_ system of vector pSEVA2514 (Aparicio *et al*., 2019b), which allows intense by short-lasting induction of the genes inserted downstream. The result of these operations were plasmids pSEVA2514-*rec2* (GenBank N° MN180223) and pSEVA2514-*rec2-mutL*_E36K_^PP^ (GenBank N° MN180222; Fig. 1). Supplementary Fig. S2B shows that this expression system overperformed the previous recombineering platform based on induction of the recombinase with 3-methyl-benzoate (3-MB) through the *xylS*-P*m* device (Ricaurte *et al*., 2018), with an order of magnitude of improvement in editing efficiency. Simultaneous expression of the recombinase and *mutL*_E36K_^PP^ with this system should therefore enable the survival of mismatches generated by recombineering within a given time window, which would otherwise be removed by an active MMR system (see below). In order to optimize the recombineering protocol, different induction times for the thermal induction of *rec2* were tested (Supplementary Fig. S2C), as little as 5 min being sufficient for achieving high levels of allelic replacements in the standard recombineering assay described by (Ricaurte *et al*., 2018).

### *Benchmarking the MMR activity of wild-type and mutS/mutL*_E36K_^PP^ *P. putida strains*

In order to obtain some reference values on the ability of MutL_E36K_^PP^ to allow inheritance of mismatches in *P. putida*, we designed two recombineering oligonucleotides (SR and NR, Fig. 2A, Table 1). These enter single-nucleotide changes that—using the *E. coli* system as an orientation—represent the extremes of the ability of the MMR system to remove mismatches. But at the same time they cause easily detectable phenotypes if incorporated in the replication fork. In one case (SR oligonucleotide; (Ricaurte *et al*., 2018)) the sequence was designed for targeting the *rpsL* gene (PP_0449) of *P. putida* EM42. This gene encodes the 30S ribosomal protein S12 and a change in the wild-type codon AAA (K43) ➔ ACA (T43) confers streptomycin resistance (Sm^R^). Upon Rec2-mediated recombineering, SR should generate an A:G mismatch predicted to show low sensitivity to the MMR system (Babic *et al*., 1996; Nyerges *et al*., 2016). If maintained, the change enters in the *rpsL* the mutation A➔C conferring Sm^R^. By the same token, oligonucleotide NR (Table 1) was designed to target the gene *gyrA* (PP_1767) of *P. putida*, (encoding a DNA gyrase subunit) for executing an amino acid change D87N known to confer resistance to nalidixic acid (Nal^R^) in *E. coli* and *P. aeruginosa* (Yoshida *et al*., 1990; Kureishi *et al*., 1994). The mismatch introduced by NR causes the same change in the *gyrA* gene of *P. putida* (GAC) D87 ➔ (AAT) N87 but in this case making two modifications at once (G➔A and C➔T). Both G:T and C:A mismatches thus ought to survive the action of MMR to result in resistance to nalidixic acid (Nal^R^). NR thus had a considerable diagnostic value, as G:T and C:A mismatches are highly sensitive to the MMR system (using again *E. coli* as provisional reference; (Nyerges *et al*., 2016).

**Figure 2.**
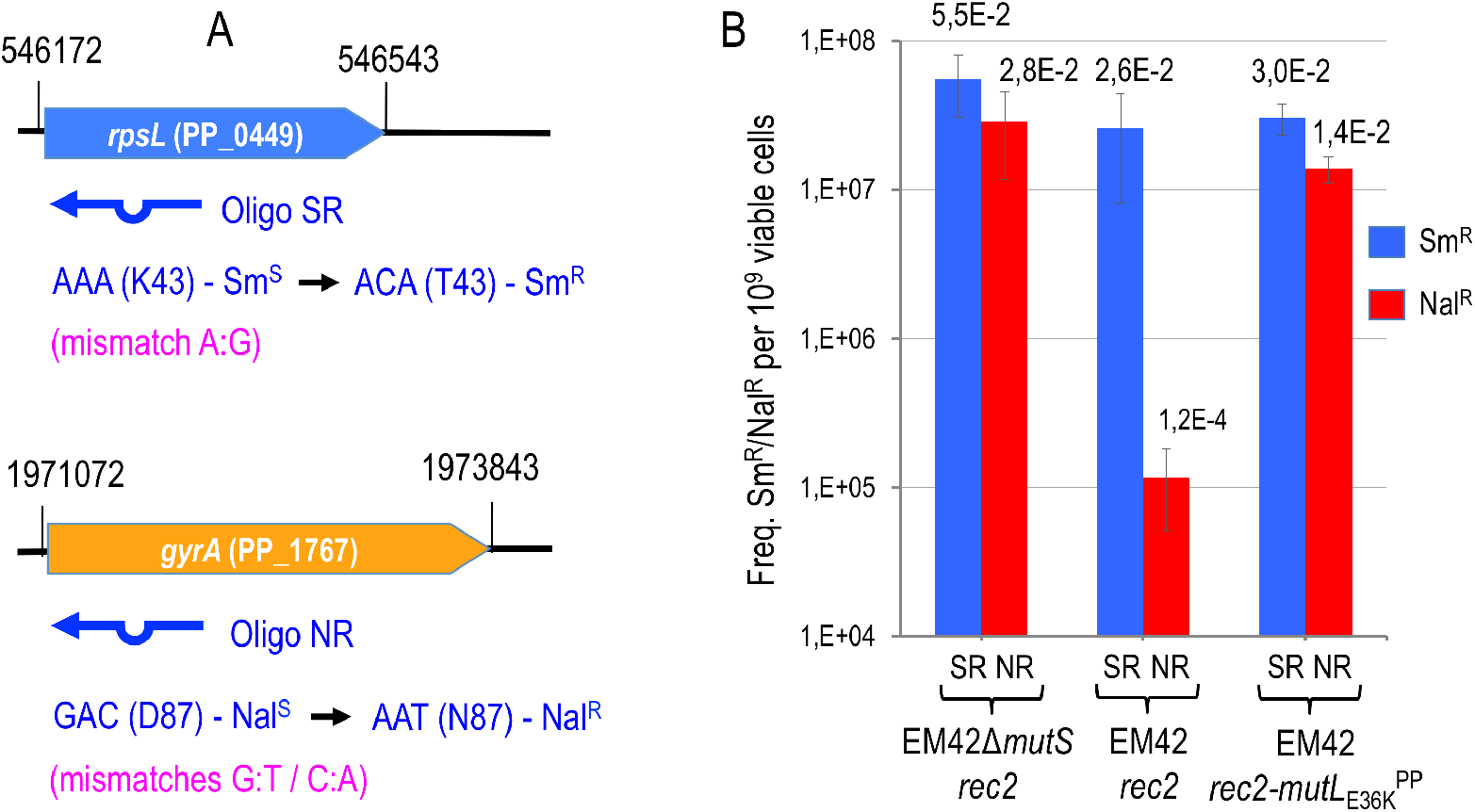
The impairment of MMR system of *P. putida* allows unbiased detection/repair of two mismatches with MMR differential sensitivity. **A.** Reporter genes used to assess MMR system activity in *P. putida* EM42 are outlined. Chromosomal coordinates and locus tag are shown. Recombineering oligonucleotides are sketched below each gene, featuring the mutation introduced, the mismatch between chromosomal and synthetic sequences and also the resulting phenotype. **B.** Oligos SR (A:G mismatch, low MMR sensitivity, confers Sm^R^) and NR (G:T and C:A mismatches, high MMR sensitivity, confer Nal^R^) were used for recombineering in *P. putida* strains Δ*mutS/*pSEVA2514-*rec2*, EM42*/*pSEVA2514-*rec2* and EM42*/*pSEVA2514-*rec2-mutL*_E36K_^PP^. Cultures of each strain were subjected to recombineering with SR and NR oligonucleotides separately as explained in Experimental Procedures section. Dilutions of each experiment were plated on LB and LB-Sm (oligo SR) or LB-Nal (oligo NR) and colonies counted after 18 h at 30 °C. Column values represent mean recombineering frequencies (mutants per 10^9^ viable cells) of two independent experiments with the standard deviation. Absolute frequencies (mutants per viable cell) are also shown above each column.

For benchmarking the experimental system to investigate the mispairing preferences of the MMR system of *P. putida*, strain EM42 was separately transformed with pSEVA2514-*rec2* and with pSEVA2514-*rec2*-*mutL*_E36K_^PP^. Also *P. putida* EM42 Δ*mutS* was transformed with pSEVA2514-*rec2*. The resulting transformants were expected to support ssDNA recombineering of the mutagenic oligonucleotides described above upon thermal activation of the P_*L*_ promoter. However, they are anticipated to have MMR in a different operational state: *P. putida* Δ*mutS* (pSEVA2514-*rec2*) has a permanently disabled system due to the deletion of the main component of MMR machinery (*mutS*); *P. putida* EM42 (pSEVA2514-*rec2*) has a wild-type MMR system; and *P. putida* EM42 (pSEVA2514-*rec2*-*mutL*_E36K_^PP^) as a functional wild-type MMR system at 30 °C which can be transiently inactivated upon thermal induction and overexpression of the dominant-negative *mutL*_E36K_^PP^ allele.

The results of the recombineering experiments run with these 3 strains upon electroporation of the SR and NR oligonucleotides are shown in Fig. 2B. In one hand, SR incorporation to the replication fork yielded Sm^R^ cells through a single nucleotide change in which the involved mismatch (A:G) is expected to be poorly recognized and thus left unrepaired in cells bearing an intact MMR machinery. On the other hand, recombineering of the NR oligonucleotide should generate Nal^R^ cells but the G:T and C:A mismatches could be readily recognized and fixed by MMR. The expected outcome of these experiments in the wild type background of *P. putida* EM42 (pSEVA2514-*rec2*) should thus show much higher recombineering efficiency with SR than with NR. In contrast, when the MMR system is impaired—whether permanently in strain *P. putida* Δ*mutS* (pSEVA2514-*rec2*) or transiently in *P. putida* EM42 (pSEVA2514-*rec2-mutL*_E36K_^PP^) the frequencies of allelic replacements using SR and NR should converge.

These predictions were not only confirmed by the data of Fig. 2B, but the results also allowed quantification of the recombineering efficiencies under the various conditions. Specifically, Fig. 2B revealed a difference of two orders of magnitude between Sm^R^ and Nal^R^ clones resulting from the recombineering experiments with the wild-type strain *P. putida* EM42 (pSEVA2514-*rec2*) and SR/NR oligos, respectively. In contrast, the frequencies of Nal^R^ resistant clones in *P. putida* Δ*mutS* (pSEVA2514-*rec2*) upon transformation with the NR oligonucleotide increased to the levels of the Sm^R^ clones of the same strain treated with the SR oligo. These data confirmed that MMR is altogether eliminated in the Δ*mutS* strain and that this lesion abolishes any bias in mispair recognition and repairing process. Finally, when the MutL_E36K_^PP^ protein was overexpressed in strain *P. putida* EM42 (pSEVA2514-*rec2-mutL*_E36K_^PP^) the frequencies of Sm^R^ and Nal^R^ resulting respectively from treatments with the SR and NR oligos were very similar. Taken together, the results of Fig. 2 showed that the inherent activity of the MMR native of *P. putida* clearly discriminates different types of mismatches (in the case tested: low activity against A:G and high activity against G:T + C:A) which can be transiently and effectively silenced *in vivo* upon expression of the dominant negative allele E36K of *mutL*. On this basis we set out to explore the whole landscape of single-nucleotide mispairings allowed or not in *P. putida*, as explained next.

### *Rationale for unraveling the hierarchy of the MMR system of* P. putida

In order to characterize the bias in the detection/repair of single nucleotide changes by the MMR system of *P. putida*, a set of four mutagenic oligonucleotides targeting the *pyrF* gene (PP_1815) of *P. putida* were designed. This gene, which is equivalent to yeast’s URA3, encodes orotidine 5’-phosphate decarboxylase and its inactivation makes cells to become resistant to fluoroorotic acid (5FOA^R^; (Galvao and de Lorenzo, 2005). The oligos for mutagenic recombineering of *pyrF* share the same sequence within the gene but bear four distinct positions fully degenerated (Supplementary Table S1, Fig. 3A), targeting nucleotides A (oligo PYR_A), C (oligo PYR_C), T (oligo PYR_T) and G (oligo PYR_G). The oligonucleotides encode also in all cases a C➔A change which turns GAA codon E58 into TAA (Stop) in the midst of the *pyrF* ORF. When incorporated into the chromosome, this change thus generates cells with a truncated, non-functional *pyrF* gene, which become then uracil auxotrophs and 5FOA^R^. This set of oligonucleotides can therefore generate all possible mismatches *in vivo* during a ssDNA recombineering experiment, thereby exposing them to the endogenous MMR activity—whether fully active, fully inactive or transiently inhibited. While all mutants that have incorporated the oligos in the genome can be selected by growing the cells in the presence of uracil and 5FOA, the frequency of the accompanying changes can be quantified by PCR and deep sequencing of the targeted region of *pyrF*.

**Figure 3.**
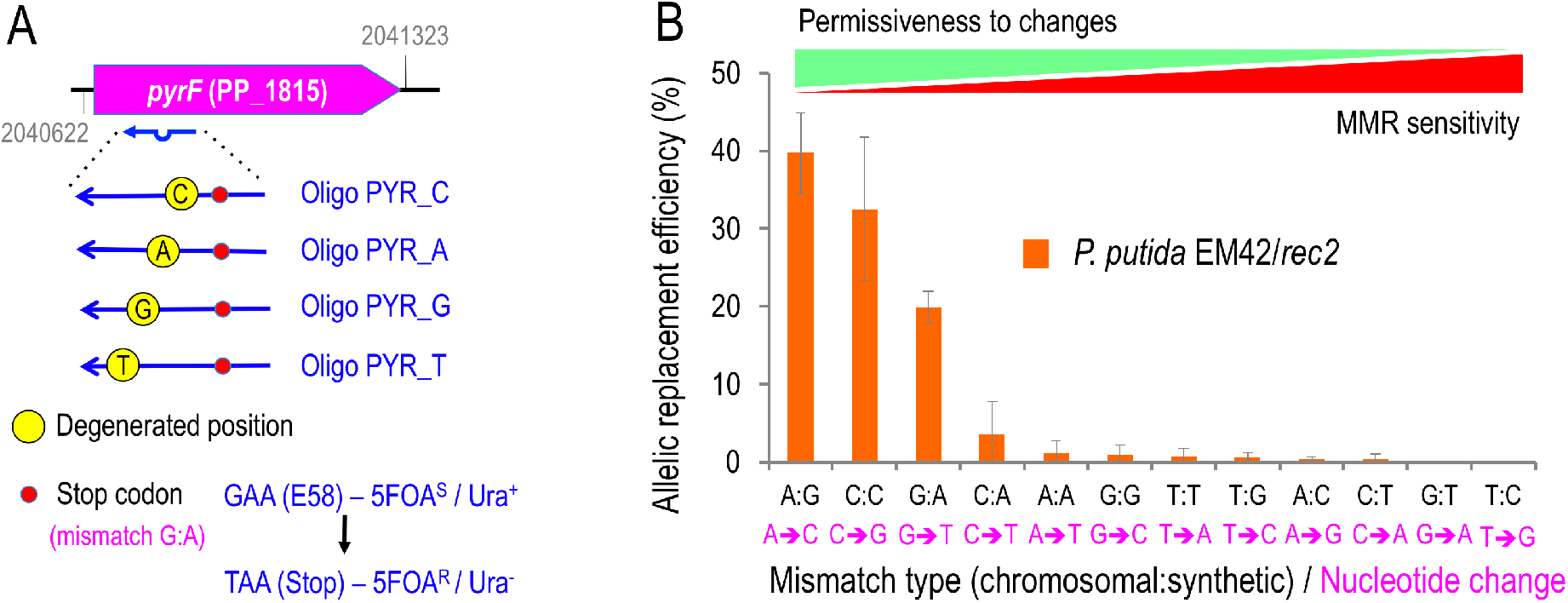
Hierarchy of *P. putida* MMR system **A.** The *pyrF* reporter gene used to assess MMR hierarchy in *P. putida* EM42 is outlined. Locus tag and chromosomal coordinates are shown. The four PYR_X oligos introduce the same Stop codon (red dot), thus rendering a *pyrF*-strain which is uracil auxotroph and 5FOA resistant, but bear a different degenerated position each (yellow dot, the genomic nucleotide that pairs with oligonucleotide sequence is depicted inside), generating three mismatches per oligonucleotide. Pictures are not drawn to scale **B.** *P. putida* EM42/pSEVA2514-*rec2* (WT strain-wild-type MMR system) was subjected to recombineering with an equimolar mixture of oligos PYR_C, PYR_A, PYR_G and PYR_T. After selection of minimal media plus Ura/5FOA, 500 *pyrF*-colonies were streaked in the same media and the streaks re-suspended in water, then pelleted and the whole genomic content extracted. *pyrF* gene was PCR amplified from the gDNA and sequenced by Illumina deep sequencing. Sequences were analysed to verify the presence of single mutations on the four degenerated positions targeted by PYR oligonucleotides. The relative frequencies of incorporated mutations were plotted and labelled with the original mismatch and the base change originated. The values are the mean of two independent experiments, bars representing standard deviations

### *Nucleotide mispairing preferences of the MMR system of* P. putida *EM42*

On the basis of the above, the predisposition of *P. putida* MMR system to recognize and repair different DNA mismatches and the ability of the MutL_E36K_^PP^ protein to effectively abrogate the bias was inspected. To this end, the oligos employed in the ssDNA recombineering experiments bear specific mismatches with the chromosomal DNA that are incorporated into the replication fork by the action of the Rec2 recombinase. Since the endogenous MMR activity can repair the mismatches to various degrees, the frequency of 5FOA^R^ mutants become a quantitative assay of MMR activity. Recombineering experiments were first performed with equimolar mixtures of PYR_A/T/C/G oligonucleotides in the wild-type, MMR^+^ background of strain *P. putida* EM42 (pSEVA2514-*rec2*) as explained in *Experimental procedures*. After selection on M9-Citrate-Ura-5FOA plates, 500 colonies were reisolated in the same medium, then pooled together and genomic DNA extracted. This DNA pool was used as the template for amplification of *pyrF* gene by PCR and the resulting amplicons were analyzed by Illumina deep sequencing (see *Experimental Procedures* for details). Fig. 3B shows the relative frequency of the allelic replacements observed in this experiment, reflecting the bias of the wild-type *P. putida* EM42 MMR system to detect and repair single nucleotide mispairings. The results allowed us to establish the following hierarchy of mismatch recognition from less to more sensitive (and thus more to less permissive to changes) : A:G< C:C< G:A< C:A, A:A, G:G, T:T, T:G, A:C, C:T< G:T, T:C. This grading is comparable, but not identical, to that found in *E. coli* (Kramer *et al*., 1984; Nyerges *et al*., 2016). Similarly to this species of reference, the MMR system of *P. putida* EM42 shows very low sensitivity to A:G and C:C mismatches (Babic *et al*., 1996; Nyerges *et al*., 2016). However, the T:T mismatch is poorly recognized/repaired in *P. putida* while the MMR system of *E. coli* has a much higher sensitivity to it. On the other hand, G:T and T:C mismatches have remarkably high repair rates in *P. putida* EM42 (i.e., less permissive), while *E. coli* do not show such noticeable differences. All in all, the MMR discrimination encompasses three orders of magnitude i.e. from A:G (39.7 % efficiency) vs. to T:C (0.02 %). According to these data, C:A/G:T, the mismatches involved in the Nal^R^ phenotype mediated by NR oligonucleotide (Fig. 2B) should be highly sensitive to MMR, but in fact the difference with the upper extreme is only two logs—surely due to the presence of the second C:A mismatch in the NR oligo. This can be explained in light of (Sawitzke *et al*., 2011), namely, if two mismatches become simultaneously incorporated, the MMR recognizes them differently from the individual mutations, a phenomenon that seems to occur also in *P. putida*. Note also that MMR action on the mismatches occurs always in the newly synthesized strand that incorporates the mutagenic oligonucleotide. As commented in the Introduction above, this hints towards a mechanism of template/newly synthesized DNA discrimination in *P. putida* which cannot depend on *dam* methylation, an open question that deserves further studies.

To clarify the role of MMR in the recognition/repair just described, the same recombineering experiments were performed with strains *P. putida* Δ*mutS* (pSEVA2514-*rec2*) and *P. putida* EM42 (pSEVA2514-*rec2-mutL*_E36K_^PP^) and the results were plotted along with those with the wild-type stain (Fig. 4A). A heatmap with the mean values of the resulting allelic replacements is shown in Fig. 4B. Both using of the MMR defective strain and heat induction of the *mutL*_E36K_^PP^ allele resulted in remarkable reduction of the recognition/repair bias of *P. putida* MMR system (Supplementary Fig. S3 shows detailed information of the allelic replacement frequencies obtained). However, inspection of the actual figures of 5FOA^R^ mutants indicated that a complete loss of MMR largely equalizes, but does not completely abolish, the bias towards stable inheritance of mutations generated by mispairings. Yet, the differential rate of repair in the MMR^+^ strain between high and low sensitive mismatches is close to 1000-fold, while the permanent or transient removal of MMR reduces the distance to not more than 4-fold (see Supplementary Fig. S3 for details). The rates of allelic replacements in mismatches very sensitive to MMR repair are particularly important in strain *P. putida* EM42 (pSEVA2514-*rec2-mutL*_E36K_^PP^) as they increase to 7% in the case of G:T (compared to a mere 0.04% in the wild-type host) and to 5% for T:C (0.02% in the MMR^+^ strain). These data not only sheds light on the recognition preference and fixing of nucleotide mismatches in *P. putida* but also accredit the dominant negative activity of MutL_E36K_^PP^ in this species and suggest general method for momentarily supressing MMR in bacteria subjected to a recombineering protocol.

**Figure 4.**
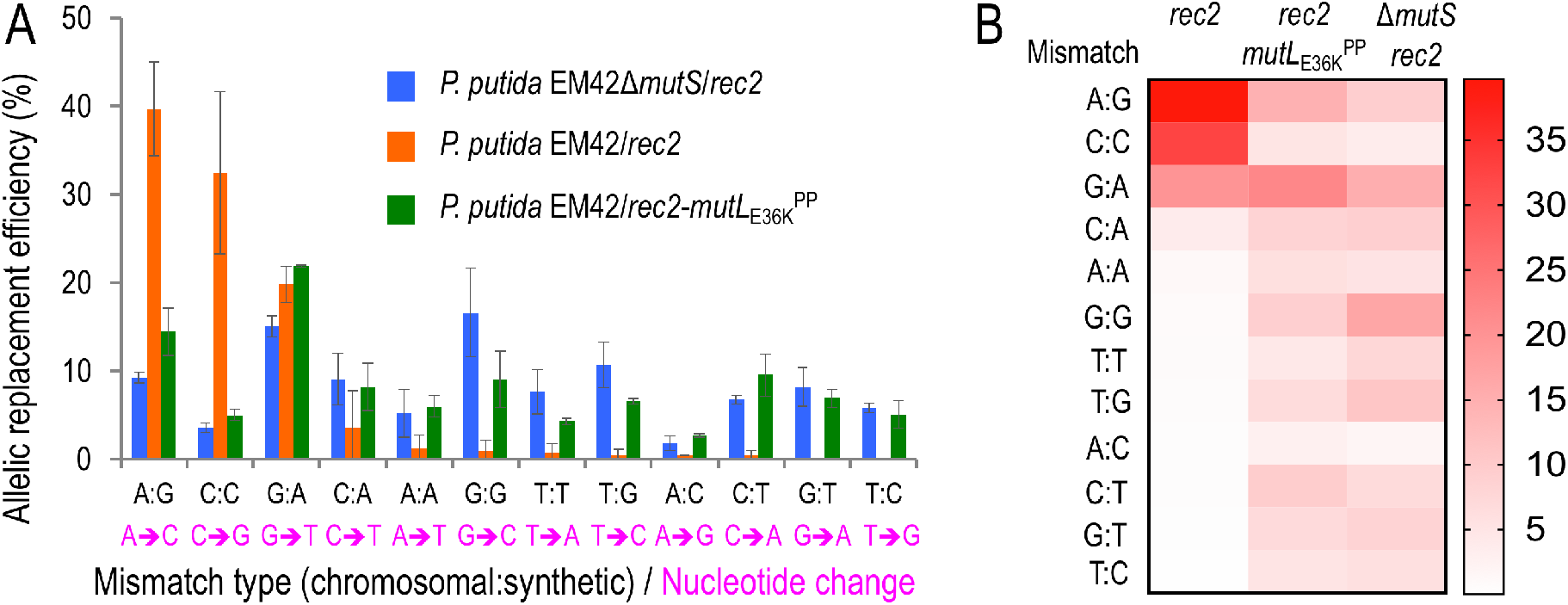
Effect of MMR inactivation on mismatch repair bias. **A.** The same experiment as shown in Fig. 3 was performed using EM42Δ*mutS/*pSEVA2514-*rec2* (Δ*mutS* strain with an inactive MMR system) and EM42*/*pSEVA2514-*rec2-mutL*_E36K_^PP^ (MMR system transiently inhibited) strains and the results were compared with the wild-type scenario to study differences in the mutation bias under constitutive or transient impairment of MMR system, respectively. **B.** Heatmap of allelic replacement frequencies of the three strains under study. Detailed information of allelic replacement frequencies of every mismatch is shown in Supplementary Fig. S3.

### Transient expression of mutL_E36K_^PP^ inhibits MMR but does not cause whole-genome mutagenesis

As indicated in Supplementary Fig. S2A, the loss of *mutS* multiplies the spontaneous mutagenesis rate of *P. putida* by at least 2 orders of magnitude as revealed with a simple count of Rif^R^ mutants in the population. One remaining question regarding the recombineering results above was whether the same could be brought about by the transient thermoinduction of *mutL*_E36K_^PP^ during the period of time involved in the recombineering experiments. This piece of information is of essence for judging whether plasmid pSEVA2514-*rec2-mutL*_E36K_^PP^ could be a useful construct for implementing specific single-nucleotide changes in the genome of *P. putida* through Rec2-mediated recombineering without the complication of suspecting off-target, concurrent mutations. To sort this out we first adopted a Rif ^R^ fluctuation-like assay for quantifying the impact of inhibiting the MMR system in the mutational rate of the strains under study. Fig. 5 shows that overexpression *mutL*_E36K_^PP^ had a minor effect on the frequency of spontaneous appearance of Rif^R^ clones of *P. putida* EM42. In contrast, the Δ*mutS* strain under the same conditions exhibited >100 fold increase in mutations leading to Rif ^R^ compared to the wild-type *P. putida*, a value in agreement with the earlier data shown in Supplementary Fig. S2A. In order to asses further the appearance of random mutations during the transient inactivation of the MMR system with *mutL*_E36K_^PP^, two Sm^R^ and two Nal^R^ colonies recovered from recombineering experiments of strain *P. putida* EM42 (pSEVA2514-*rec2-mutL*_E36K_^PP^) with SR and NR oligonucleotides, respectively, were submitted to whole genome sequencing. In addition, a Sm^R^ clone originated in the treatment of the control *P. putida* EM42 (pSEVA2514-*rec2*) strain expressing only Rec2 was also analyzed. Genomic DNA of each of the 5 strains was purified and sequenced by Illumina. As shown in Supplementary Table S2, zero to three SNPs were detected in strains transiently expressing *mutL*_E36K_^PP^, while no mutations could be observed in the control MMR^+^ strain. This demonstrated a very low level of spontaneous mutations when transiently inhibiting the MMR system of *P. putida*—low enough to consider thermoinduction of *mutL*_E36K_^PP^ an ideal asset for implementing high-efficiency genome editing methods in *P. putida*.

**Figure 5.**
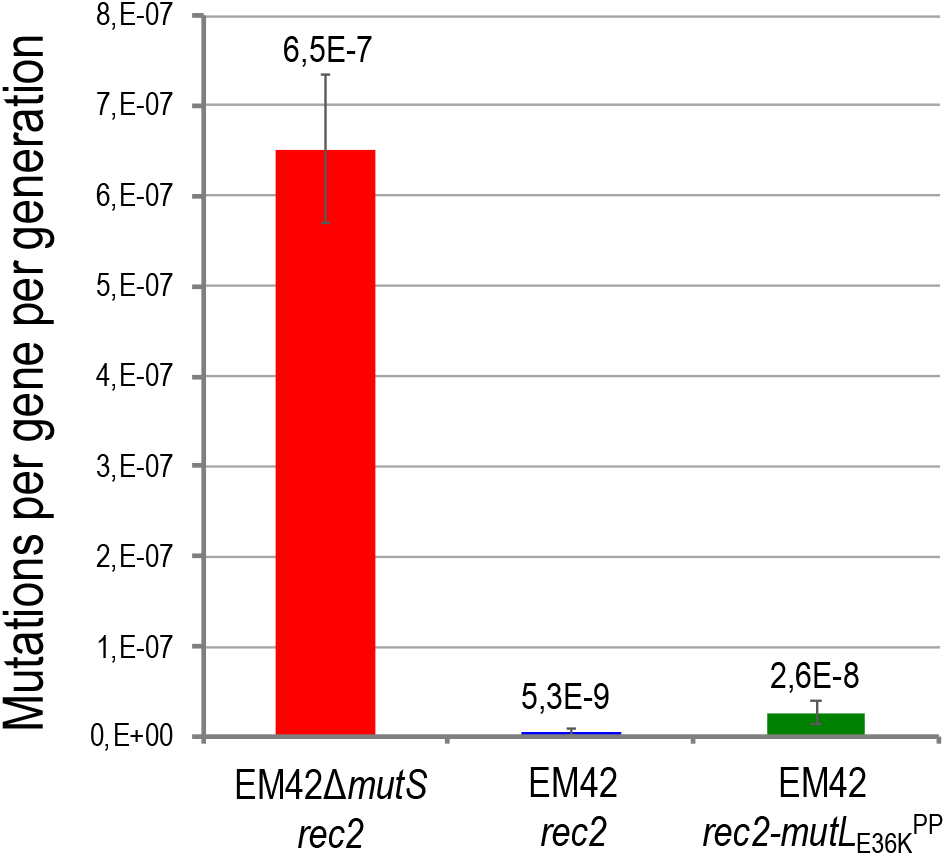
Mutation rates of *P. putida* EM42-derived strains. A rifampicin resistance fluctuation assay was used to estimate the mutation rates of EM42Δ*mutS/*pSEVA2514-*rec2*, EM42*/*pSEVA2514-*rec2* and EM42*/*pSEVA2514-*rec2-mutL*_E36K_^PP^ as described in Experimental Procedures. Fifteen independent replicas were performed and results analyzed with the FALCOR web tool by the MMS-Maximum Likelihood Estimator Method. FALCOR averages estimating mutations per gen per generation are depicted above the columns. Error bars account for the 95% Confidence Intervals difference.

### Conclusion

The results of the experiments described above provide three valuable pieces of information on how *P. putida* handles mismatches in the DNA helix of its genomic DNA caused by replication errors, misincorporation of damaged bases or (as exploited in this work) deliberate mispairings artificially introduced by means of recombineering strategies. The advantage of the last is that one can recapitulate all possible changes in a controlled fashion, as we have done in the present work. **First,** *P. putida* has an active MMR system that includes at least the *mutS* and *mutL* homologs of *E. coli* and brings about a distinct hierarchy of mismatch recognition and suppression. But possibly, the MMR system of *P. putida* (as is the case with *P. aeruginosa* (Oliver *et al*., 2002)) comprises other activities as well: permanent or transient suppression of these genes dramatically reduce, but does not entirely eliminate the bias in tolerance to different types of mismatches. **Second,** as in any MMR device known in other bacteria, the system discriminates the old template strand of DNA from the newly synthesized sequence that bears the mismatched nucleotide. How this happens in the absence of *dam* methylation is unknown, although similarly to the case of *Bacillus* it could involve the beta clamp of the replication machinery (Simmons *et al*., 2008). And **third,** transient expression of the dominant negative E36K of *mutL* along with the *rec2* recombinase (any possibly any other ssDNA recombinase active in this species) creates a window of opportunity for introduction of all types of chromosomal single-base changes without significant offsite mutations. This property, empowered by plasmid pSEVA2514-*rec2-mutL*_E36K_^PP^ (Fig. 1) will enable expansion of advanced methods of recombineering-based genome engineering such as DIvERGE (Nyerges *et al*., 2018) or pORTMAGE-based technology (Nyerges *et al*., 2016) towards this environmentally and biotechnologically important bacterium.

## EXPERIMENTAL PROCEDURES

### Strains, media and general procedures

Liquid LB was used as routine growth media (10 g l^−1^ tryptone, 5 g l^−1^ yeast extract, and 5 g l^−1^ NaCl) for *E. coli* and *P. putida* strains used in this study (Table 1). Glycerol-free Terrific Broth (TB; 12 g l^−1^ tryptone, 24 g l^−1^ yeast extract, 2 g l^−1^ KH_2_PO_4_, 9.4 g l^−1^ K_2_HPO_4_) was used for after-electroporation recovery during recombineering experiments. Bacterial strains were cultivated with shaking (170 rpm) at 30 °C (*P. putida*) or 37 °C (*E. coli*). M9 minimal media (Sambrook *et al*., 1989) was supplemented, when stated, with 0.2% w/v citrate for *P. putida* growth. Solid media was prepared adding 15 g/L of agar to liquid media. When necessary, liquid and solid media were supplemented with 50 μg ml^−1^ of kanamycin (Km), 15 μg ml^−1^ of gentamicin (Gm), 30 μg ml^−1^ of chloramphenicol (Cm), 100 μg ml^−1^ of streptomycin (Sm), 100 μg ml^−1^ rifampicin (Rif), 50 μg ml^−1^ of nalidixic acid (Nal), 20 μg ml^−1^ of Uracil (Ura) or 250 μg ml^−1^ of 5-fluoroorotic acid (5-FOA). Standard DNA manipulations were conducted according to manufacturer recommendations and previously established protocols (Sambrook *et al*., 1989). Gibson Assembly was carried out as outlined in (Aparicio *et al*., 2017) using a home-made reaction mixture. Genomic DNA was isolated with the DNAeasy^®^ UltraClean^®^ Microbial Kit (Qiagen). Following manufacturer recommendations, PCR amplifications for cloning purposes were performed with Q5 polymerase (New England Biolabs) while DNA Amplitools Master Mix was used for diagnosis PCRs. Plasmids were introduced in *P. putida* strains via tripartite mating as described in (Martinez-Garcia and de Lorenzo, 2012). Oligonucleotides were purchased from Sigma with the exception of PYR_C, PYR_A, PYR_G, PYR_T and all PCR primers used for *pyrF* deep sequencing, which were synthesized at the Nucleic Acid Synthesis Laboratory of the Biological Research Centre, Hungarian Academy of Sciences, Szeged (Hungary) and purified using high-performance liquid chromatography (HPLC). Primers were suspended in 1× Tris-EDTA (TE) buffer (pH 8.0) at 100 μM final concentration.

### Plasmid construction

To obtain pSEVA2514-*rec2*, in which the *rec2* recombinase is under the control of the heat-inducible *c*I857-P_*L*_ expression system, the *xylS*-P*m* induction module of pSEVA258-*rec2* was substituted with the thermo-inducible expression module of pSEVA2514. Both plasmids were restricted with PacI/AvrII and the 6.0 Kb of band of pSEVA258-*rec2* was ligated to the 1.4 Kb band of pSEVA2514 and the ligation transformed in *E. coli* CC118. For the construction of pSEVA2514-*rec2-mutL*_E36K_^PP^, in which *mutL*_E36K_^PP^ allele is co-expressed with *rec2*, the corresponding sequence was first assembled in the pSEVA258-*ssr* and later on transferred to pSEVA2514-*rec2*. pSEVA258-*ssr-mutL*_E36K_^PP^ was generated as follows: first, *mutL* of *P. putida* KT2440 (2.0 Kb) was colony amplified with oligos mutL-KT-Fw/mutL-KT-Rv (Tm= 65 °C, 1 min. elongation, Q5 polymerase). This PCR fragment was Gibson assembled with BamHI/SphI restricted pSEVA258-*ssr* and the assembly mix transformed in *E. coli* CC118. The resulting pSEVA258-*ssr-mutL* was used as a template for amplifying two fragments: a 0.9 Kb fragment containing the 3’-end of *ssr* and the 5’-end of *mutL* with Gibson-PP-Beta-Fw/ mutLKT-Gibson-2 (Tm= 60 °C, 30 sec. elongation, Q5 polymerase) and a fragment containing a 0.4 Kb segment of *mutL* with mutLKT-Gibson-3/ mutLKT-Gibson-4 (Tm= 71 °C, 30 sec. elongation, Q5 polymerase). mutLKT-Gibson-2 and mutLKT-Gibson-3 primers incorporate the single nucleotide change responsible of the amino acid change E36→K36 in the *mutL* ORF to generate *mutL_E36K_. pSEVA258-ssr-mutL* was cut with EcoRI and the 8.4 Kb fragment, containing the 5’-end of *ssr* and the 3’-end of *mutL*, was Gibson assembled with the two PCR fragments described above to eventually generate pSEVA258-*ssr-mutL*_E36K_^PP^. Finally, this plasmid was used as a template for amplifying *mutL*_E36K_ with primers mutL_E36K_-Gib-Fw/ mutL_E36K_-Gib-Rv (Tm= 65 °C, 1 min. Elongation, Q5 polymerase) and the resulting 2.0 Kb fragment was Gibson assembled with pSEVA2514-*rec2* restricted with XbaI/HindIII. The assembly mix was transformed in *E. coli* CC118 to obtain pSEVA2514-*rec2-mutL*_E36K_^PP^. pSEVA231-C-mutS1 bears a CRISPR array with a spacer targeting the *mutS* gene of *P. putida* KT2440. Spacer design and cloning was performed as described previously (Aparicio *et al*., 2018, 2019a). Briefly, oligonucleotides cr-mutS-S-1 and cr-mutS-AS-1 were annealed and the resulting dsDNA fragment was ligated into pSEVA231-CRISPR restricted with BsaI. Ligation was transformed in *E. coli* CC118. All plasmids constructed in this work, either by Gibson Assembly or by restriction/ligation, were transformed in *E. coli* CC118 calcium chloride competent cells, selected in LB-Km solid media and colonies checked by miniprep+restriction. Inserts were fully sequenced (Macrogen Spain) to verify the accuracy of the constructs.

### *Construction of* P. putida *EM42* ΔmutS *strain by recombineering/CRISPR-Cas9*

The deletion protocol described in (Aparicio *et al*., 2018) was applied on *P. putida* EM42 bearing pSEVA658-*ssr* and pSEVA421-Cas9tr plasmids. Recombineering with MAGE-mutS-2 oligonucleotide and CRISPR-Cas9 counterselection with pSEVA231-C-mutS1 plasmid was used to delete a 0.7 Kb segment of the *mutS* gene (PP_1626) of *P. putida* EM42. One out of fifty colonies checked by PCR with primers mutS-check3/ mutS-check4 (Tm= 55 °C, 30 seconds elongation) yielded the 0.6 Kb fragment expected for the deletion mutant.

### Rifampicin Assay

The mutational rate of *P. putida* EM42 Δ*mutS* and its parental strain (*P. putida* EM42) was estimated by monitoring the appearance of rifampicin resistant (Rif ^R^) colonies. Overnight cultures grown in LB were adjusted to OD_600_ ≈ 1.0 and 1 ml (~10^9^ cells) of each sample was centrifuged 1 minute at 11.000 rpm. The pellets were re-suspended in 100 μl of LB and plated on LB-Rif solid media. Rif ^R^ colonies were counted after 24 hours of incubation at 30 °C. Two independent replicas were done and the medias and standard deviations were represented as the frequencies of Rif ^R^ mutants per 10^9^ cells.

### ssDNA recombineering protocol mediated by thermal induction

Recombineering experiments were accomplished basically as described in (Ricaurte *et al*., 2018). Some modifications were implemented to trigger the activation of the thermo-inducible *c*I857-P_*L*_ expression system of pSEVA2514 derivatives driving the expression of *rec2* and *mutL*_E36K_^PP^ genes. Overnight cultures of *P. putida* EM42 bearing the plasmids under study were back-diluted to OD_600_= 0.1 in a total volume of 20 ml of LB-Km in 100 ml Erlenmeyer flasks. Cultures were incubated at 30 °C/ 170 rpm until OD_600_= 0.4-0.5. Then, flasks were transferred to a water bath at 42 °C for 5 minutes with gentle shaking to increase quickly the temperature and induce the expression of Rec2/ MutL_E36K_^PP^ proteins. Flasks were incubated 10 additional minutes at 42 °C in an air shaker at 250 rpm (15 minutes of total induction at 42 °C) and then cooled down in ice for 5 minutes to stop the thermal induction. When stated, different induction times were applied with shorter or longer incubations in the air shaker. Competent cells were prepared at RT by centrifuging cultures at 3220 g/ 5 minutes and washing the pellets three consecutive times with 10, 5 and 1 ml of 300 mM sucrose solution. Pellets were finally resuspended in 200 μl of the same solution. One hundred microliters of this suspension were added with 1 μl of the recombineering oligonucleotide (stock at 100 mM), mixed thoroughly and the mixture transferred to an electroporation 2 mm-gap width cuvette (Bio-Rad). Electrotransformation was performed in a Micropulser™ device (Bio-Rad Laboratories, Hercules, CA, USA) at 2.5 kV and cultures were immediately inoculated in 5 ml of fresh TB and recovered overnight at 30 °C/ 170 rpm. Several dilutions of the recovered cultures were plated in the appropriate selective and non-selective solid media depending on the current experiment (see below).

### ssDNA recombineering experiments with SR and NR oligonucleotides

Recombineering with oligonucleotides SR (A:G mismatch, low MMR sensitivity; A→C change produces a Sm-resistant phenotype) and NR (G:T and C:A mismatches, low and high MMR sensitivity, respectively; G→A and C→T changes produce a Nal-resistant phenotype) was performed as described in the previous section on strains *P. putida* EM42/ pSEVA2514-*rec2* (WT-wild type MMR system), *P. putida* EM42Δ*mutS*/ pSEVA2514-*rec2* (Δ*mutS-* inactive MMR system) and *P. putida* EM42*/* pSEVA2514-*rec2-mutL*_E36K_^PP^ (transient MMR inactivation upon expression of *mutL*_E36K_^PP^ protein). After overnight recovery, dilutions 10^−2^, 10^−3^, 10^−4^ and 10^−5^ of SR and NR electroporated cultures were plated on LB-Sm and LB-Nal, respectively, to select cells harbouring the allelic replacements. To estimate the number of viable cells, dilutions 10^−7^ and 10^−8^ of were plated on LB. Plates were incubated 18 hours at 30 °C and absolute colony counts were taken. The recombineering frequency (RF) was calculated as the ratio between the number of antibiotic-resistant colonies and the number of viable cells. This ratio was normalized to 10^9^ viable cells. In order to check the accuracy of the allelic replacements, ten clones from each strain/oligonucleotide experiment were PCR amplified either for *rpsL* gene (Sm-resistant colonies coming from SR experiments-oligos rpsL-Fw/ rpsL-Rv, Tm= 57 °C, 45 seconds elongation, 0.8 Kb product) or for *gyrA* gene (Nal-resistant colonies from NR experiments-oligos gyrA-Fw/ gyrA-Rv, Tm= 57 °C, 45 seconds elongation, 0.4 Kb product). PCRs were purified and sequenced with rpsl-Fw and gyrA-Fw, respectively (Macrogen Spain). All 60 clones analyzed showed the expected changes introduced by the recombineering oligonucleotides without additional mutations in the region sequenced.

### MMR recognition hierarchy in P. putida EM42

For a more detailed characterization of the MMR system of *P. putida* EM42, the three strains studied above were subjected to recombineering with a mixture of oligonucleotides PYR_A, PYR_C, PYR_T and PYR_G. 10 μl of each oligonucleotide at 100 mM were mixed and 2 μl of the mixture (0.5 μl of each oligonucleotide) were used for electrotransformation. The ssDNA recombineering protocol mediated by thermal induction was applied as explained before but cultures were allowed to recover only 5 hours at 30 °C/ 170 rpm since longer recovery times in *pyrF*-targeted experiments were reported to decrease the appearance of *pyrF*-mutants (Ricaurte *et al*., 2018). After recovery, several dilutions of each culture (10^−2^, 10^−3^) were plated on M9-Citrate-Ura-5FOA and plates were incubated 48 hours at 30 °C to allow the slow-growing *pyrF*-colonies to appear. Five hundred colonies were streaked in the same media and incubated as before. The 500 streaks were pooled together by suspension in 2 ml of water, the sample centrifuged 1 minute at 11.000 rpm and the pellets used for genomic DNA (gDNA) extraction with DNeasy^®^ UltraClean^®^ Microbial Kit (Qiagen). A negative control experiment was also performed with *P. putida* EM42*/* pSEVA2514-*rec2* electroporated without any oligonucleotide. In this case, the post-electroporated culture was recovered 5 hours and plated in M9-Citrate-Ura-5FOA (10^−1^ and 10^−2^ dilutions) and M9-Citrate (10^−5^ and 10^−6^ dilutions). As expected, only few colonies appeared on the selective media, all of them showing the fast-growing phenotype typical of spontaneous mutants, non-*pyrF* related, described elsewhere (Galvao and de Lorenzo, 2005; Aparicio *et al*., 2016). A pool of approximately 10.000 colonies rescued from the M9-Citrate plates were used for gDNA extraction and served as a control of the downstream process of deep amplicon sequencing of *pyrF*. An independent replica of this set of four experiments was performed and gDNA samples from both replicas were used to estimate the activity of the MMR system by deep amplicon sequencing of the *pyrF* targeted region.

### Deep amplicon sequencing of pyrF

To determine the allelic composition of *pyrF* at the oligonucleotide-target site, we utilized a previously described Illumina MiSeq deep sequencing protocol (Nyerges *et al*., 2016). To create Illumina sequencing libraries, a 138 nucleotide-long region of *pyrF* in *P. putida* that was targeted by recombineering-oligonucleotides (PYR_G, PYR _A, PYR_T and PYR _C), was PCR amplified from the previously isolated, pooled gDNA samples using the corresponding barcoded primer pairs specifically designed for each experiment/sample (Supplementary Table S1). To multiplex sequencing samples on Illumina MiSeq, barcoded PCR primers were designed based on a previously published protocol (Kozich *et al*., 2013) and consisted of the appropriate Illumina adaptor sequences, a 10 nucleotide-long pad sequence, and a 2 nucleotide-long linker besides the terminal genomic target-specific primer sequences. Besides barcoded PCR primers, custom Illumina sequencing primers according to (Kozich *et al*., 2013) were also designed (Supplementary Table S1). Next, the *pyrF* oligo-target region from each gDNA sample was amplified in 4×25 μl volumes, consisting of 50 μl 2× Q5 Hot-Start MasterMix (New England Biolabs), 2 μl of the corresponding sample-specific, barcoded, reverse Illumina primer (100 μM) plus 2 μl PYR_ILMF (100 μM) primer, 200 ng template gDNA and 45 μl nuclease-free H2O. PCRs were performed in thin-wall PCR tubes in a BioRad CFX96 qPCR machine with the following thermal profile: 98 °C 3 minutes, 23 cycles of (98 °C 15 seconds, 62 °C 20 seconds, 72 °C 20 seconds) and a final extension of 72 °C for 5 minutes. Following PCRs, the 180 basepair-long amplicons were purified by using a Zymo Research DNA Clean and Concentrator™ Kit according to the manufacturer’s protocol (Zymo Research) and eluted in in 30 μl 1× Tris-EDTA (TE) buffer (pH 8.0). To prepare samples for sequencing, amplicons were quantified using Qubit dsDNA BR assay kit (Thermo Fisher Scientific), mixed, and libraries were sequenced on an Illumina MiSeq instrument using v2 paired-end 2×250-cycle sequencing kit (Illumina). To perform sequencing, the Illumina MiSeq cartridges were supplemented with 100 μM stocks of our custom Illumina sequencing primers (Supplementary Table S1). After sequencing, raw sequencing reads were de-multiplexed according to their corresponding barcodes. The average sequencing read counts were 160000 per sample. Next, the overlapping read-pairs were identified and merged to yield one template-read from each combined sequencing read using pandaseq v2.8 (Masella *et al*., 2012). Reads were then trimmed to an error probability threshold of 0.001 (Phred quality = 30) using readtools 1.5.2 (Gomez-Sanchez and Schlotterer, 2018). Merged paired-end reads were then mapped to their corresponding reference sequence (*P. putida pyrF-*PP_1815) by using bowtie2 2.3.4 (Langmead) in “--very-sensitive-local” mode and the nucleotide composition was extracted for each nucleotide position within the oligo-targeted region. Allelic replacement frequencies at each oligo-targeted nucleotide positions were quantified by measuring the distribution and ratio of nucleotide substitutions for each reference nucleotide position (Nyerges *et al*., 2016). Finally, the allelic replacement frequency of each individual substitution was normalized to the sum of all substitutions detected in the experiment. Data from two independent replicas of each experimental condition were used to calculate medias and standard deviations.

### Mutational rate measurement by a fluctuation-like assay

A rifampicin resistance fluctuation assay was performed with *P. putida* EM42 (pSEVA2514-*rec2*) *P. putida* Δ*mutS* (pSEVA2514-*rec2*) and *P. putida* EM42 (pSEVA2514-*rec2-mutL*_E36K_^PP^). The strains were inoculated in 3 ml of LB-Km and incubated overnight at 30 °C/170 rpm. In order to mimic the experimental conditions of a standard recombineering experiment, overnight cultures were back-diluted to OD_600_ ~ 0,1 in 3 ml fresh LB-Km and incubated at 30 °C/170 rpm until OD_600_ ~ 0,5. Cultures were then placed in a water bath at 42 °C for 5 minutes with gentle shaking, transferred to an air shaker at 42 °C/ 250 rpm/ 10 min (total incubation at 42 °C= 15 min) and incubated at 4 °C for 5 min. After overnight growth at 30 °C/170 rpm aliquots of each culture were plated on LB and LB-Rif and plates incubated 24 hours at 30 °C. Total colony count was done and the data from fifteen independent replicas of the experiment were used to calculate the mutational rate of each strain by using the Ma-Sandri-Sarkar Maximum Likelihood Estimator method and the FALCOR web tool (Hall *et al*., 2009).

### Whole genome sequencing and bioinformatics for SNPs detection

Genomic DNA samples of *P. putida* EM42 query strains were sequenced in Macrogen Inc. (Korea). Truseq PCR Free Libraries of 350 bp were processed in Illumina Hiseq2500 (2×100 bp) flow cells (output coverage ~900x). Quality of raw data was analyzed using FASTQ files with FastQC tool (https://www.bioinformatics.babraham.ac.uk/projects/fastqc/). No quality issues were detected and Illumina reads were aligned to *P. putida* KT2440 genome (NC 002947.4) using “bwa aln” and “bwa sampe” commands with default parameters (Li and Durbin, 2010). Alignment files in SAM format were compressed, coordinate-sorted and indexed using “samtools view -bS”, “samtools sort” and “samtools index” commands, respectively (Li *et al*., 2009). Before coordinate-sorted step, duplicated reads (paired reads aligning exactly at the same genomic coordinates, considered as PCR artifacts) were removed with “samtools rmdup” command. Since the genome of *P. putida* EM42 contains 10 deletions of variable size compared with the reference genome of *P. putida* KT2440, genomic regions with no coverage were detected in SAM/BAM files using “bedtools genomecov -bga” (Quinlan and Hall, 2010) and parsing the output with “grep -w 0$”. SNP detection was carried-out using “samtools mpileup -B” and “bcftools call - m” (Li, 2011). Biological impact of detected polymorphisms was determined with snpEff tool setting upstream and downstream gene regions 500 bp in size (-upDownStreamLen 500;Cingolani *et al*., 2012). Only variants with QUAL > 200 and coverage (DP) > 200 were considered for SNP validation.

## ACKNOWLEDGEMENTS

This work was funded by the HELIOS Project of the Spanish Ministry of Science BIO 2015-66960-C3-2-R (MINECO/FEDER); the ARISYS (ERC-2012-ADG-322797), MADONNA (H2020-FET-OPEN-RIA-2017-1-766975), BioRoboost (H2020-NMBP-BIO-CSA-2018), and SYNBIO4FLAV (H2020-NMBP/0500) Contracts of the European Union and the S2017/BMD-3691 InGEMICS-CM funded by the Comunidad de Madrid (European Structural and Investment Funds). CS was supported by grants from the European Research Council H2020-ERC-2014-CoG 648364, the Wellcome Trust, GINOP-2.3.2-15-2016-00020, GINOP-2.3.2-15-2016-00014 and the Lendület Program of the Hungarian Academy of Sciences. AN was supported by a PhD fellowship from the Boehringer Ingelheim Fonds.

