## Supplementary information for "Mismatch repair hierarchy of *Pseudomonas putida* revealed by mutagenic ssDNA recombineering of the *pyrF* gene"

SUPPLEMENTARY INFORMATION to Aparicio et al.

**Figure S1.** Alignment of MutL proteins of *E. coli* and *P. putida* KT2440.

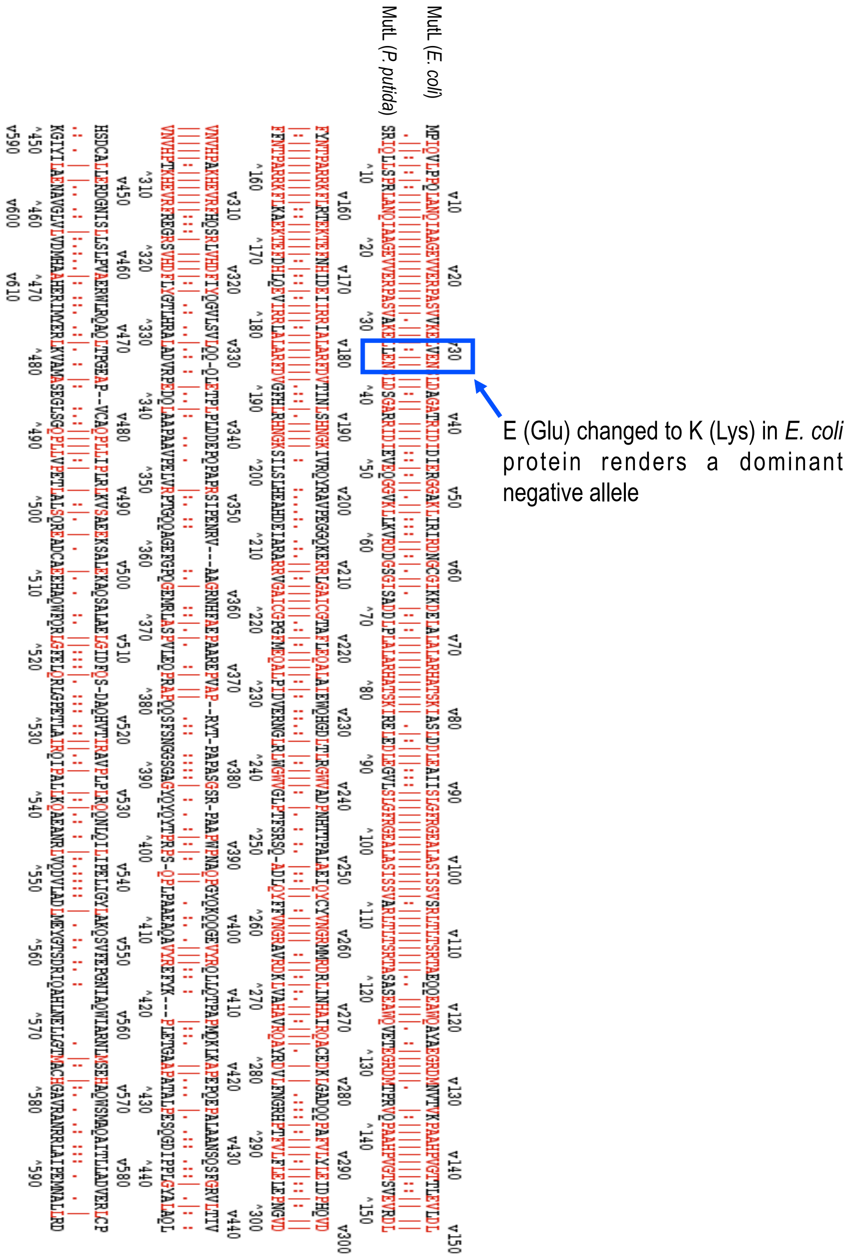


A square and an arrow show the amino acid position ($E_{32}$) responsible of the dominant-negative phenotype of allele *mutLE32K* of *E. coli* (Nyerges et al., 2016). This position, along with the first half of the protein, is well conserved in the *P. putida* homologue.

**Figure S2**. Mutational activity of *P. putida* EM42 Δ*mutS* strain and recombineering performance of Rec2 recombinase under the *c*I8567-$P_L$ expression system

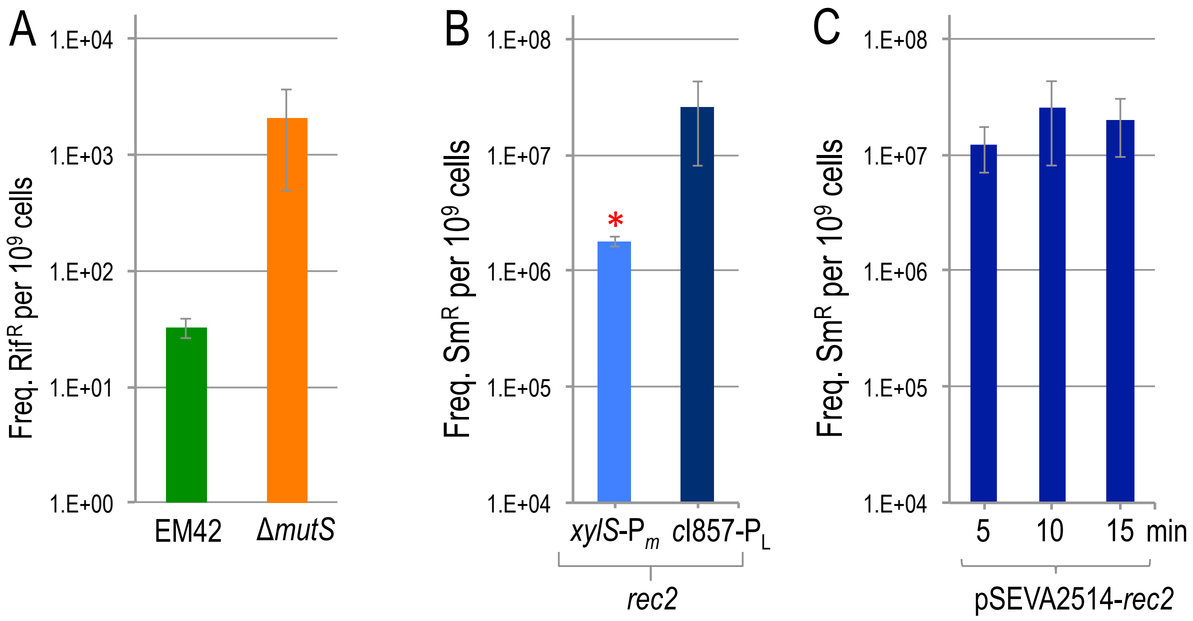


**A.** As a proxy of the mutational activity of *P. putida* EM42 Δ*mutS* and its parental strain, a Rifampicin assay was developed. Results are shown as the number of Rif resistant colonies per $10^9$ cells **B.** Comparison of editing efficiencies using SR oligonucleotide in *P. putida* EM42 strains harbouring *rec2* under either *xylS*-$P_m$ (pSEVA258-*rec2*) or *c*I857-$P_L$ (pSEVA2514-*rec2*) expression systems. Respectively, 30 minutes of induction with 3-MB 1 mM and 30 minutes of thermal-shock at 42 ºC were applied in standard recombineering experiments (see Experimental Procedures for details). Dilutions of each experiment were plated on LB and LB-Sm and colonies counted after 18 h at 30 ºC. The values are the means of two independent experiments, bars representing standard deviations **C.** Comparison of editing efficiencies of Rec2 upon thermal induction for 5, 10 and 15 minutes at 42 ºC using SR oligonucleotide. Experiments were performed as described above. *Recombineering data for *xylS*-$P_m$-*rec2* reproduced from (Ricaurte et al., 2018)

**Figure S3.** Allelic replacement frequencies mediated by recombineering with PYR oligonucleotides. All mismatches are expressed chromosomal: synthetic.

| | | Allelic replacement frequencies (%) | | |
|---|---|---|---|---|
| **Change** | **Mismatch** | ***P. putida* EM42 ∆mutS *rec2*** | ***P. putida* EM42 *rec2*** | ***P. putida* EM42 *rec2-mutLE36K*** |
| A ➔ C | A:G | 9,24 ± 0,61 | 39,7 ± 5,25 | 14,50 ± 2,66 |
| C ➔ G | C:C | 3,58 ± 0,61 | 32,49 ± 9,19 | 4,99 ± 0,61 |
| G ➔ T | G:A | 15,08 ± 1,19 | 19,85 ± 2,10 | 21,91 ± 0,13 |
| C ➔ T | C:A | 9,08 ± 2,91 | 3,64 ± 4,09 | 8,20 ± 2,65 |
| A ➔ T | A:A | 5,22 ± 2,69 | 1,17 ± 1,55 | 5,97 ± 1,25 |
| G ➔ C | G:G | 16,62 ± 4,97 | 0,91 ± 1,24 | 9,10 ± 3,23 |
| T ➔ A | T:T | 7,66 ± 2,46 | 0,73 ± 0,99 | 4,31 ± 0,37 |
| T ➔ C | T:G | 10,76 ± 2,55 | 0,50 ± 0,61 | 6,58 ± 0,33 |
| A ➔ G | A:C | 1,84 ± 0,85 | 0,48 ± 0,08 | 2,76 ± 0,18 |
| C ➔ A | C:T | 6,8 ± 0,49 | 0,42 ± 0,55 | 9,56 ± 2,36 |
| G ➔ A | G:T | 8,22 ± 2,23 | 0,04 ± 0,00 | 6,94 ± 1,02 |
| T ➔ G | T:C | 5,84 ± 0,52 | 0,02 ± 0,03 | 5,11 ± 1,60 |

**Supplementary Table S1.** Oligonucleotides used in this study.

| Name | Sequence (5' → 3')[a] | Usage |
|---|---|---|
| SR | G*T*C*A*GACGCACACGGCATACTTTACGCAGTG CCGAGTTAGGTTTTGTCGGCGTGGTGGTGTACAC ACGGGTGCACACGCCACGACGCTGC | Recombineering oligo for *rpsL* gene: AAA (K43) changed to ACA (T43), mismatch A:G, confers Sm resistance |
| NR | AACGAGAACGGCTGGGCCATACGCACGATGGTA TTGTAGACCGCAGTGTCGCCGTGCGGGTGGTA | Recombineering oligo for *gyrA* gene: GAC (D87) changed to AAT (N87), mismatches G:T and C:A, confers Nal resistance |
| PYR_C | C*T*TGAGGTCCAGGAACACTTAGAAGCCCTTGTC ACACAGNGTTTCGACAATGCCCGAAGCGCTGCTG GTGAACAGCTCCTTGCCAACC*T*T | Recombineering oligo for *pyrF* gene: "A" changes GAA (E58) into TAA (stop codon), mismacth G:A; "N" stands for a degenerated position, mismatch C:N |
| PYR_A | C*T*TGAGGTCCAGGAACACTTAGAAGCCCTTGTC ACACAGGGNTTCGACAATGCCCGAAGCGCTGCT GGTGAACAGCTCCTTGCCAACC*T*T | Recombineering oligo for *pyrF* gene: "A" changes GAA (E58) into TAA (stop codon), mismacth G:A; "N" stands for a degenerated position, mismatch A:N |
| PYR_G | C*T*TGAGGTCCAGGAACACTTAGAAGCCCTTGTC ACACAGGGTTTNGACAATGCCCGAAGCGCTGCT GGTGAACAGCTCCTTGCCAACC*T*T | Recombineering oligo for *pyrF* gene: "A" changes GAA (E58) into TAA (stop codon), mismacth G:A; "N" stands for a degenerated position, mismatch G:N |
| PYR_T | C*T*TGAGGTCCAGGAACACTTAGAAGCCCTTGTC ACACAGGGTTTCGNCAATGCCCGAAGCGCTGCT GGTGAACAGCTCCTTGCCAACC*T*T | Recombineering oligo for *pyrF* gene: "A" changes GAA (E58) into TAA (stop codon), mismacth G:A; "N" stands for a degenerated position, mismatch T:N |
| rpsL-Fw | GACATGAAATGTTGCCGATG | With rpsL-Rv, to amplify part of *rpsL* gene of *P. putida* (0.8 Kb) |
| rpsL-Rv | CTGTTCTTGCGTGCTTTGAC | With rpsL-Fw, to amplify part of *rpsL* gene of *P. putida* (0.8 Kb) |
| gyrA-F | GGCCAAAGAAATCCTCCCGGTCAA | With gyrA-R, to amplify part of *gyrA* gene of *P. putida* (0.5 Kb) |
| gyrA-R | AGCAGGTTGGGAATACGCGTCG | With gyrA-F, to amplify part of *gyrA* gene of *P. putida* (0.5 Kb) |
| mutL-KT-Fw | CCTCACCAAGTCGGCCAGCCGGCCGCAGGAGTA AGGATCCAGAAGGAGAATATACCATGAGTGGCG GTTCGCG | With mutl-KT-Rv, to amplify *mutL* gene of *P. putida* KT2440 |
| mutL-KT-Rv | AGGGTTTTCCCAGTCACGACGCGGCCGCAAGCT TGCATGCTCATCGACCGCGCAGGA | With mutl-KT-Fw, to amplify *mutL* gene of *P. putida* KT2440 |
| Gibson-PP-Beta-Fw | TGGAGTCATGACCATGCCTAGGCCGCGGCCGCG CGAATTCAGAAGGAGAATATACCATGAGCCAAGT AGCCAGGGTC | With mutLKT-Gibson-2,to amplify *ssr* and $N_t$ part of *mutL* |
| mutLKT-Gibson-2 | CACCGGAGTCCAGGCTGTTTTTCAGCAGTTCCTT GGCCAC | With Gibson-PP-Beta-Fw, to amplify *ssr* and $N_t$ part of *mutL*, It includes the change E36→K36 to generate mutLE36K |
| mutLKT-Gibson-3 | GTGGCCAAGGAACTGCTGAAAAACAGCCTGGAC TCCGGTG | With mutLKT-Gibson-4, to amplify the $C_t$ part of *mutL*. It includes the change E36→K36 to generate $mutL_{E36K}^{PP}$ |

| | | |
|---|---|---|
| mutLKT-Gibson-4 | AGCGCCAGCCGCCGGATC | With mutLKT-Gibson-3, to amplify the $C_t$ part of *mutL* |
| mutLE36K-Gib-Fw | GGAGCCCGTTGATGCAAATCATCACTGAGGATCCTCTAGAAGAAGGAGAATATACCATGAGTGGCGGTTCGCG | With mutLE36K-Gib-Rv, to amplify $mutL_{E36K}^{PP}$ |
| mutLE36K-Gib-Rv | GTCGCCAGGGTTTTCCCAGTCACGACGCGGCCGCAAGCTTTCATCGACCGCGCAGGA | With mutLE36K-Gib-Fw, to amplify $mutL_{E36K}^{PP}$ |
| MAGE-mutS-2 | T*T*C*G*CGACCAAGGACCTCAAGGGCTTCGGCTGCGACAAGCTGACCAACCGCGTGCACGGCTACTTCATCGAACTGCCGACCAAGCAGGCC | Recombineering oligo to delete a 0,7 Kb fragment of *mutS* gene of *P. putida* EM42 |
| mutS-check3 | GGCGGTACTGGACATCAC | With mutS- check3,to amplify part of *mutS* gene of *P. putida* |
| mutS-check4 | AGTTCAGGTCGAGGTTCAG | With mutS-check3,to amplify part of *mutS* gene of *P. putida* |
| cr-mutS-S-1 | 5´p-AAACCCGGCTATGACAACGAACTGGACGAGCTGCG | Oligo to obtain the *mutS1* spacer |
| cr-mutS-AS-1 | 5´p-AAAACGCAGCTCGTCCAGTTCGTTGTCATAGCCGG | Oligo to obtain the *mutS1* spacer |
| PYR26R | CAAGCAGAAGACGGCATACGAGATGCTCATCGGAATATTATTTGGTGTTGGGAATGTCGTGGA | To amplify *pyrF in P. putida* EM42Δ*mutS*/ pSEVA2514-*rec2* Sample 1 |
| PYR28R | CAAGCAGAAGACGGCATACGAGATCTTTTGCGGAATATTATTTGGTGTTGGGAATGTCGTGGA | To amplify *pyrF* in *P. putida* EM42Δ*mutS*/ pSEVA2514-*rec2* Sample 2 |
| PYR29R | CAAGCAGAAGACGGCATACGAGATTAGTTGCGGAATATTATTTGGTGTTGGGAATGTCGTGGA | To amplify *pyrF* in *P. putida* EM42/ pSEVA2514-*rec2* Sample 1 |
| PYR31R | CAAGCAGAAGACGGCATACGAGATATCGTGCGGAATATTATTTGGTGTTGGGAATGTCGTGGA | To amplify *pyrF* in *P. putida* EM42/ pSEVA254-*rec2* Sample 2 |
| PYR32R | CAAGCAGAAGACGGCATACGAGATTGAGTGCGGAATATTATTTGGTGTTGGGAATGTCGTGGA | To amplify *pyrF* in *P. putida* EM42/ pSEVA2514-*rec2*, no oligo (CONTROL) Sample 1 |
| PYR33R | CAAGCAGAAGACGGCATACGAGATCGCCTGCGGAATATTATTTGGTGTTGGGAATGTCGTGGA | To amplify *pyrF in P. putida* EM42/ pSEVA2514-*rec2*, no oligo (CONTROL) Sample 2 |
| PYR34R | CAAGCAGAAGACGGCATACGAGATGCCATGCGGAATATTATTTGGTGTTGGGAATGTCGTGGA | To amplify *pyrF* in *P. putida* EM42/ pSEVA254-*rec2*- $mutL_{E36K}^{PP}$ Sample 1 |
| PYR35R | CAAGCAGAAGACGGCATACGAGATAAAATGCGGAATATTATTTGGTGTTGGGAATGTCGTGGA | To amplify *pyrF* in *P. putida* EM42/ pSEVA2514-*rec2*- $mutL_{E36K}^{PP}$ Sample 2 |
| PYR_ILMF | AATGATACGGCGACCACCGAGATCTACACATCGTATATGGTAATTTATGGATTTCCCTACCCGTGAGG | Forward primer to amplify *pyrF* with barcoded PYRXXR primers |
| Illumina-Fw seq primer | CGTATATGGTAATTTATGGATTTCCCTACCCGTGAGGC | For Illumina deep sequencing of pyrF |
| Illumina-Rv seq primer | CGGAATATTATTTGGTGTTGGGAATGTCGTGGAACTTGAG | For Illumina deep sequencing of pyrF |
| Illumina-barcode seq primer | CTCAAGTTCCACGACATTCCCAACACCAAATAATATTCCG | For Illumina deep sequencing of pyrF |

Asterisks denote phosphorothioate linkages. 5´ hemi-sequence of deletion oligonucleotide MAGE-mutS-2 is shown in blue color. Single changes introduced by recombineering oligonucleotides and mutL-KT-Fw/Rv primers are highlighted in red. 5´ phosphorilated oligonucleotides are depicted as 5´p-. The Shine-Dalgarno sequence and the start codon of *ssr*, *mutL* and *mutL*$_{E36K}^{PP}$ genes are shown underlined in Gibson-PP-Beta-Fw, mutL-KT-Fw and mutLE36K-Gib-Fw primers. PYRXXR primers were used for deep sequencing of pyrF gene and reverse-complement sequence of barcodes are shown in orange.

**Supplementary Table S2.** Off-target mutations during transient inhibition of MMR System.

| *P. putida* EM42 | Selected phenotype | Nº SNPs | Locus | Change | Position (NB_002947.4) | Quality/DP |
|---|---|---|---|---|---|---|
| pSEVA2514-*rec2* | Sm$^{R}$ | 0 | - | - | - | - |
| pSEVA2514-*rec2*-*mutL*$_{E36K}^{PP}$ | Sm$^{R}$-Col1 | 0 | - | - | - | - |
| pSEVA2514-*rec2*-*mutL*$_{E36K}^{PP}$ | Sm$^{R}$-Col2 | 3 | PP_0158 | G→A | 169.482 | 225/519 |
| | | | PP_5547 | C→T | 3.383.400 | 225/649 |
| | | | PP_5075 | T→C | 5.792.744 | 225/582 |
| pSEVA2514-*rec2*-*mutL*$_{E36K}^{PP}$ | Nal$^{R}$-Col6 | 0 | - | - | - | - |
| pSEVA2514-*rec2*-*mutL*$_{E36K}^{PP}$ | Nal$^{R}$-Col7 | 3 | PP_1355 | T→C | 1.545.111 | 225/417 |
| | | | PP_3316 | T→C | 3.753.418 | 228/518 |
| | | | PP_4947 | G→A | 5.631.097 | 228/571 |
